# Stress-induced endogenous-dsRNAs drive beta-cells to IFN-I state

**DOI:** 10.1101/2023.11.10.566600

**Authors:** Kevin K. Chi, Parijat Senapati, Amy Leung, Dustin E. Schones

**Affiliations:** Department of Diabetes Complications and Metabolism, Arthur Riggs Diabetes & Metabolism Research Institute, City of Hope, Duarte, CA, 91010; Irell and Manella Graduate School of Biological Sciences, City of Hope, Duarte, CA 91010

**Keywords:** *Alu*s, dsRNAs, epigenetics, inflammatory stress, retrotransposons, viral mimicry

## Abstract

**Aims/hypothesis:** Endogenous dsRNAs originating from retrotransposons within the human genome have the potential to trigger autoimmunogenic recognition via RIG-like receptors such as MDA5. This phenomenon, known as “viral mimicry”, induces potent interferon-beta expression, resulting in cell death and potential collateral death to surrounding cells. Viral mimicry has been mostly studied in relation to epigenetic therapeutics, although there are indications the phenomenon is relevant in other physiological contexts. Here, we demonstrate that proinflammatory cytokine stress leads to the dysregulation of retrotransposons in human beta-cells, resulting in beta-cell death via viral mimicry.

**Methods:** Human islets from donors were treated with/without proinflammatory cytokines (IFN-γ, TNF-α, IL-1β) and immunostained for dsRNA. Publicly available transcriptome and chromatin accessibility data from proinflammatory cytokine treated EndoC-βH1 cells were analyzed to examine changes in transcriptional regulatory activity at retrotransposon loci. Transcriptome profiling experiments in 1.1B4 and MIN6 cells were performed to examine the generality of the results. Adenosine to inosine (A-to-I) editing was compared between sample groups to gauge the extent of RNA editing on dsRNAs. Additional transcriptome data from normal vs. type 1 diabetic human islets was also assessed for differences in A-to-I editing activity between sample groups.

**Results:** Proinflammatory cytokine stress increases the amount of dsRNA in human islets, co-incident with cells that express insulin. In the EndoC-βH1 human beta-cell line, proinflammatory cytokine stress leads to a substantial increase in transcription at RTE loci, including SINE/*Alu* elements and LINE elements, along with increased expression of RLRs (including MDA5), inflammatory transcription factors (e.g. IRF3, STAT1), and most critically, IFNB1, a potent instigator of type 1 interferon responses. Similar trends were observed in 1.1B4 and MIN6 cells. Gene ontology analysis revealed an enrichment of genes related to interferon signaling and inflammation. Inflammatory stress furthermore leads to an increase in chromatin accessibility at many RTE loci, with enrichment in a variety of RTE subfamilies (including L1, *Alu*Y, *Alu*Sp, *Alu*Sq2, and *Alu*Sc elements). Binding sites for inflammatory transcription factors, such as IRFs, are also enriched at these loci. While inflammatory stress increased the overall A-to-I editing events at *Alu* elements, the pool of total unedited *Alu*s was increased upon inflammatory stress. Examination of A-to-I editing in RNA-seq data from type 1 diabetic human islets and normal controls revealed that editing in diabetic islets is dramatically lower (∼1%) than islets from normal controls (∼15%).

**Conclusions/interpretation:** Transcription and chromatin accessibility are both increased at RTEs (notably *Alu*s) upon proinflammatory cytokine stress, accompanied by an increase in RLR expression and a shift to a type I interferon state, as indicated by increased IFNB1 and other interferon-stimulated genes, including MDA5, an RLR gene which is also a type 1 diabetes risk gene. The overall increase in A-to-I editing upon inflammatory stress is associated with increases in ADAR1 expression, yet the increase in overall unedited *Alu*s shows that cytokine stress leads to a higher dsRNA burden in the beta-cell. This increase in dsRNA burden is further compounded by increased MDA5 expression, exacerbating the inflammatory state of the beta-cell. Comparisons of A-to-I editing between normal vs. type 1 diabetic human islets suggest that type 1 diabetics have more unedited dsRNAs, hindering their ability to prevent autoimmunogenic recognition.

**Research in context:** *What is already known about this subject?*

- Cytokine stress during insulitis is a feature of autoimmune diabetes.
- Stress-induced dysregulation of retrotransposons can lead to a dsRNA-based type-I interferon response in a variety of cells (‘viral mimicry’)
- Endogenous dsRNAs are subject to adenine to inosine (A-to-I) editing by ADAR1 to prevent self-recognition.

*What is the key question?*

- Can cytokine stress drive beta-cells into an IFN-I state via recognition of endogenous dsRNAs derived from retrotransposons that lose silencing under cytokine stress?

*What are the new findings?*

- Cytokine stress causes an increase in dsRNAs derived from retrotransposons in beta-cells from human islets.
- The SINE/Alu subset of these endogenous dsRNAs are less edited in stressed beta-cells, making them more self-recognizable.
- Type 1 diabetic human islets exhibit more unedited *Alu*s.

*How might this impact on clinical practice in the foreseeable future?*

- Innate autoimmunity via endogenous dsRNAs can cause damage to beta-cells; therapies to target this pathway may help treat autoimmune diabetes.

## INTRODUCTION

Retrotransposons are viral-like genomic sequences that are silenced through transcriptionally repressive mechanisms including DNA methylation. Dysregulation of retrotransposons can lead to widespread aberrant transcription of many elements [1–4]. A subset of these transposons have the ability to generate double-stranded RNAs (dsRNAs) which can act as ‘viral mimics’ recognized by intracellular RIG-like receptors (RLRs). This can induce a potent type-I interferon (IFN-I) response, leading to cell dysfunction and death [4–6]. This ‘viral mimicry’ phenomenon can be summarized as an *inappropriate* autoimmune recognition of endogenous retroviral elements as exogenous viruses, leading to a deleterious and prolonged IFN-I response in the cell through the action of RLRs.

Inflammatory stress of the pancreatic islet (“insulitis”) occurs in the pre-diagnostic stages of diabetes during the onset of beta-cell dysfunction and death, which becomes chronic and exacerbated as the disease progresses [7–9]. Proinflammatory cytokines, including IFN-γ, TNF-α, and IL-1β, act as mediators of insulitis that can lead to beta cell dysfunction and even cell death [10, 11]. Though transcriptome and chromatin profiling have unveiled key regulatory pathways in the beta-cell inflammatory response [10–13], our understanding of this process remains incomplete. For example, the heterogeneous nature of insulitis and autoimmunity (i.e., not all beta-cells are attacked simultaneously) [13, 14] is not well understood.

Given the known involvement of retrotransposons in inflammatory processes [15–19], we hypothesized that viral mimicry is involved in insulitis and its role in type 1 diabetes pathogenesis. The rationale for our investigation into dsRNA-mediated viral mimicry in insulitis stems from a variety of precedent research. Within discordant type 1 diabetic-monozygotic twins, hypervariable DNA-methylation of immune effector cells was found to be a defining feature in the diabetic twin, suggesting a connection between epigenetic dysregulation and diabetes [20]. Individuals with type 1 diabetes have been observed to have increased IFN-I signaling (as gauged by *IFNB1* expression), suggesting activation of RLRs [9, 13, 21]. Additionally, landmark GWAS studies on type 1 diabetic individuals have implicated one such RLR, *MDA5*/*IFIH1,* an antiviral dsRNA-surveillance protein for IFN-I signaling, as a top risk locus for disease onset [11, 22]. Gain-of-function (e.g. A946T) variants of MDA5 have been shown to contribute to a higher risk of type 1 diabetes, while loss-of-function (e.g. I923V) variants contribute to lower risk [12, 22, 23]. Studies in NOD mice, a model of type 1 diabetes with a high incidence of diabetes arising from beta-cell destruction, revealed that both single- and double-knockout of *Mda5* protected the mice from occurrence of diabetes [24]. These data indicate that endogenous factors may be triggering MDA5 and guiding beta-cells towards the autoimmune IFN-I phenotype typically observed in type 1 diabetics.

Studies of MDA5 mutant structure and function reveal that this dsRNA sensor can recognize endogenous dsRNAs formed from retrotransposon loci, including human ERV and *Alu* elements [4, 6]. In normal cells, endogenous dsRNAs undergo A-to-I editing catalyzed by ADAR1, an RNA deaminase, preventing inappropriate recognition of endogenous (“self”) dsRNAs by introducing “bubbles’’ into the secondary structure of the dsRNAs, sterically hindering the RNA-helicase domain of RLRs like MDA5 and preventing attachment to edited dsRNAs [5, 25–28]. The most abundant class of dsRNAs targeted by ADAR1 are inverted repeat *Alu*s (IR-*Alu*s) [6, 29]. These arise from the transcription of two *Alu* elements proximal and inverted with respect to each other, which subsequently anneal into a dsRNA hairpin loop. ADAR1 knockdown studies demonstrate its importance in preventing endogenous dsRNAs (such as IR-*Alu*s) from triggering downstream innate autoimmune pathways [4, 6, 26].. The abundance of both ADAR1 and MDA5 and the abundance of intracellular dsRNA all likely contribute to the delicate balance between self- and non-self-recognition and aberrant activation of IFN-I pathways.

Despite the previous research suggesting a role for a viral mimicry like phenomenon triggered by insulitis to contribute to beta-cell death, a director role for viral mimicry in type 1 diabetes has not been explored. We set out to investigate the hypothesis that insulitis can act as a stressor to provoke the generation of dsRNAs from repetitive elements, thus inducing beta-cells into an IFN-I state, leading to loss of cell function/viability.

## RESULTS

### Proinflammatory cytokine-induced transcription at retrotransposons results in dsRNA generation

To test the potential that dsRNAs are upregulated in human islets in response to proinflammatory cytokine stress, we examined the abundance of dsRNAs in human islets treated with or without inflammatory cytokine stress (TNF-α and IFNγ) for 24 hours. Using an antibody that targets a 40bp dsRNA-secondary structure epitope (J2), we quantified dsRNA levels in control and treated cells with immunofluorescence. We found an increase in the abundance of dsRNAs upon 24h of proinflammatory cytokine treatment in normal human islets (**Fig. 1a,b**). Analogous experiments were carried out in 1.1B4 cells and similar trends were observed (**ESM** **Fig. 1**). Co-immunostaining with an antibody targeting insulin demonstrated that the punctate dsRNA signals are coincident with insulin presence. Quantification of the dsRNA signal indicated that beta-cells, defined by insulin presence, have a significant increase of dsRNAs upon proinflammatory stress (*p* < 0.05; Student’s *t*-test).

**Fig. 1.**
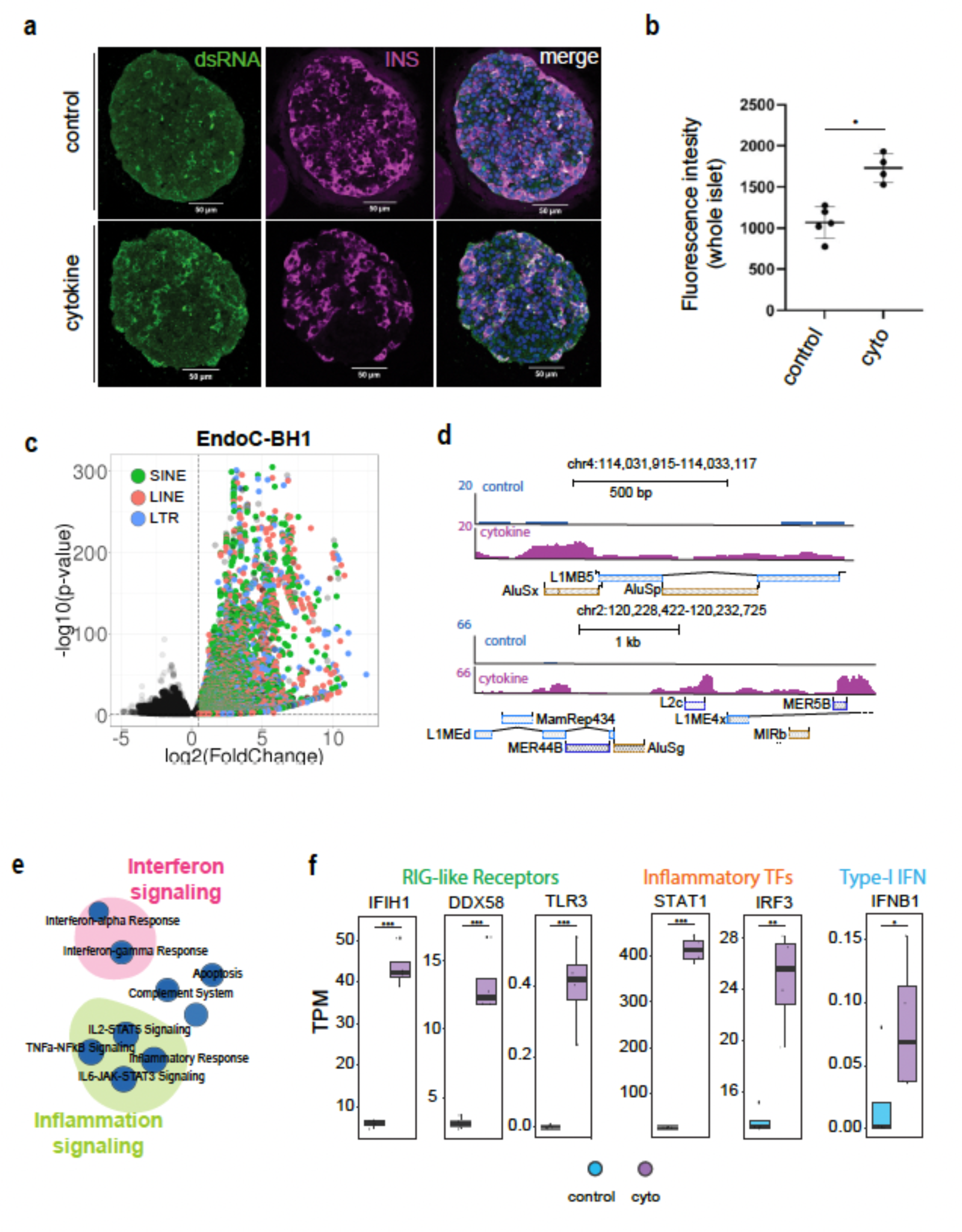
dsRNAs are increased upon proinflammatory cytokine stress. (**a**) Immunofluorescence of human islets staining for dsRNAs (green) and insulin (magenta), with DAPI staining for nuclei (blue). White scale bar represents 50um. (**b**) dsRNA fluorescence was normalized and quantified using Fiji/ImageJ. (**c**) Volcano plot of RNA-sequencing from untreated (n = 5) vs. cytokine-treated (IFN-γ + IL-1β) EndoC-βH1 (n = 5). Each point represents an annotated retrotransposon, as denoted by RepeatMasker (hg38). (**d**) Representative view of the RNA-seq signal at two example intergenic loci, showing the characteristics of cytokine-induced increase of expression at retrotransposons. (**e**) GSEA pathway analysis of EndoC-βH1 RNA-seq reveals the most enriched gene pathways upon inflammatory stress. Size of circles represent a higher proportion of enriched genes in the gene pathway. The more well-defined “hallmark gene sets” and “curated gene sets” available from MSigDB were used. (**f**) Expression levels of RLRs and interferon-related genes (* = *p* < 0.05, ** = *p* < 0.01, *** = *p* < 0.001; Student’s t-test). TPM obtained from untreated (n = 5) and cytokine-treated (n = 5) EndoC-βH1 RNA-seq.

To explore the role that retrotransposon activity has in response to proinflammatory stress, we examined publicly available EndoC-βH1 RNA-seq data from Ramos-Rodriguez and colleagues [10], including untreated (n = 5) and cytokine-treated (IL-1β + IFNγ ; n = 5) sample groups. Focusing on RepeatMasker [30] annotated retrotransposable element (RTE) loci (*hg38* build), we observed that cytokine treatment led to an increase in transcripts at RTE regions of the genome (*p* < 0.05, Wald test, Benjamini-Hochberg corrected; log2(fold-change) > 1; see Methods for details), contrasting with the much less prominent set of suppressed RTEs. (**Fig. 1c**). The ability of some RTEs to be transcribed and form dsRNAs is the main driving force in the viral mimicry phenotype – these dsRNA-capable RTEs are pervasive and may include elements from the LTR/ERV, L1/LINE, and *Alu*/SINE families [2, 3]. A closer look at two representative loci on the volcano plot shows the RNA-seq profiles of the cytokine-responsive RTEs residing at such loci in EndoC-βH1 cells. Significant increases of transcription were seen at a variety of intergenic RTEs, suggesting a pronounced dysregulation of these RTEs during the inflammatory response (**Fig. 1d**). Intriguingly, we were able to see regions where transcription increased at *Alu*/SINE and L1/LINE elements, suggesting that cytokine stress can potentially lead to problematic transcription of endogenous dsRNAs within the cell. GSEA analysis showed other transcriptional changes in beta-cells, with enrichment in “Interferon Signaling” and “Inflammation Signaling” pathways (**Fig. 1e**). In addition, we detected a significant increase in the expression of dsRNA-sensitive RLRs such as IFIH1/MDA5, DDX58/RIG-I, TLR3, as well as inflammatory transcription factors (TFs) such as IRF3 and STAT1 (*p* < 0.05; ncon = 5, ncyto = 5; Student’s *t*-test). Critically, expression of the potent type I interferon IFNB1 was increased upon cytokine stress (*p* < 0.05; ncon = 5, ncyto = 5; Student’s *t*-test) (**Fig. 1f**). 1.1B4 cells treated with inflammatory cytokines similarly showed upregulation of retrotransposon loci (**ESM** **Fig. 2**). To test for the potential for this phenomenon to exist in mouse cells as well, MIN6 cells were treated with TNF-α and IFN-γ for 24 hours. Immunofluorescence was used to examine dsRNA levels in control and treated cells and RNA-seq was performed to examine the transcription of retrotransposons. As with the EndoC-βH1 and 1.1B4 cells, MIN6 cells showed increased dsRNAs and transcription of repetitive elements upon inflammatory cytokine treatment (**ESM** **Fig. 3**). In summary, these data show that cytokine-stressed human islets exhibit an increase in dsRNA signal that tends to localize around insulin-competent cells and that cytokine treatment induces a prominent number of RTEs to be transcribed, with potential to form dsRNA. We also observed concurrent increases in dsRNA-sensing RLRs.

**Fig. 2.**
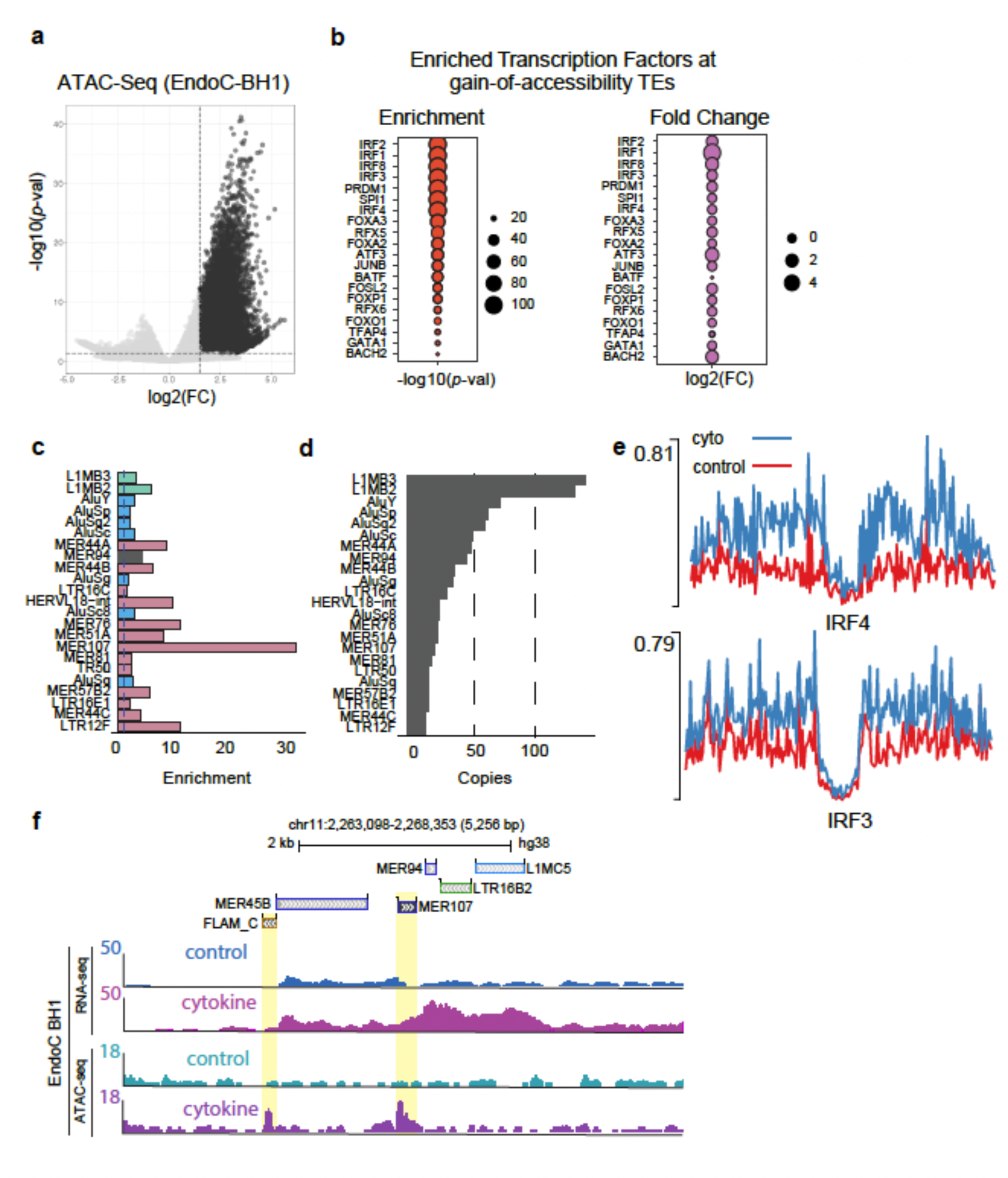
Chromatin accessibility is enriched at retrotransposon loci upon inflammatory stress. (**a**) Volcano plot of ATAC-sequencing from untreated (n = 5) and cytokine-treated (IFN-γ + IL-1β) EndoC-βH1 (n = 5). Each point represents an annotated retrotransposon, as denoted by RepeatMasker (hg38). Upper-right quadrant represents “gain-of-accessibility loci”. (**b**) Gain-of-accessibility loci were assessed for transcription factors binding sites. (left) Significance of the transcription factor, as determined by HOMER. (right) Fold-change expression of transcription factors. (**c**) Enrichment score of the gain-of-accessibility loci. Dotted blue line represents the significance threshold for enrichment of retrotransposon. Colors represent retrotransposon family (green = LINE, blue = SINE, pink = ERV). (**d**) A selection of gain-of-accessibility loci that were found to have the highest copy number to gain accessibility upon cytokine treatment. (**e**) Footprint plots showing the bias-corrected normalized signal at IRF4 and IRF3 binding sites. (**f**) Two example loci show overlap of gain-of-accessibility peaks and transcriptional upregulation at an intergenic locus, a FLAM_C/*Alu* and MER107 element.

**Fig. 3.**
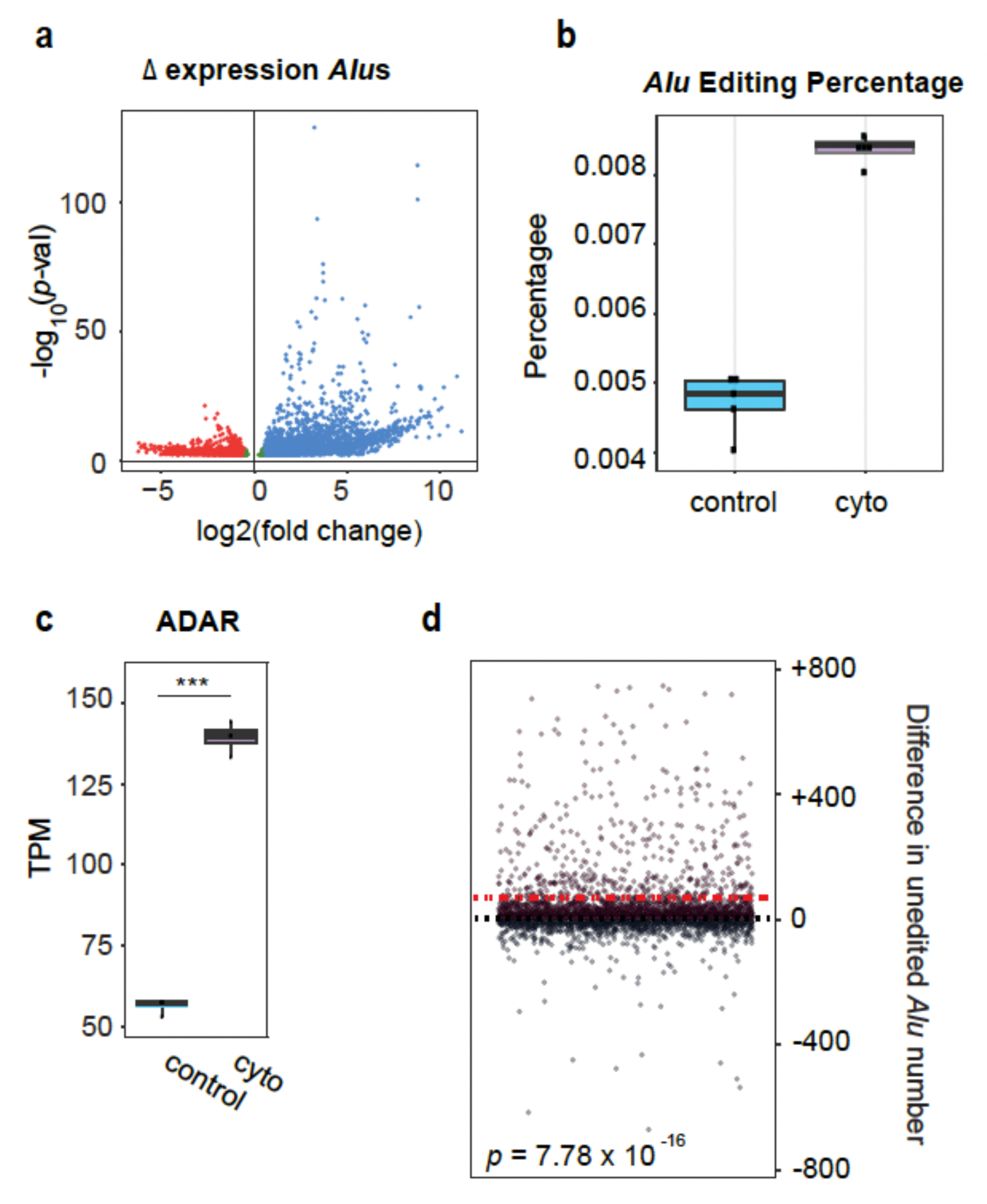
Unedited *Alu* elements increase upon inflammatory stress in human islets. (**a**) Volcano plot of RNA-sequencing from untreated (n = 5) and cytokine-treated (n = 5) human islets. Each point represents an annotated *Alu* element, as denoted by RepeatMasker (hg38). (**b**) The percentage of *Alu* editing seen in both untreated and cytokine-treated sample groups. (**c**) Expression (in TPM) of ADAR1 between untreated and cytokine-treated sample groups (*p* < 0.001; Student’s t-test). (**d**) Dot-plot showing the number of unedited *Alu*s when comparing between untreated and cytokine-treated human islets. Dots above the dashed line indicate *Alu* elements that showed less A-to-I editing instances (i.e. *Alu*s that exhibited less editing) in the cytokine-treated group compared to the untreated group. Dots below the dashed line indicate the opposite, where an *Alu* element showed more A-to-I editing instances (i.e. fewer unedited *Alu*s) in the cytokine-treated group compared to the untreated group. Red dotted-line represents the mean of all points (*p* < 7.78 x 10^16^; Kolmogrov-Smirnov test).

Using chromatin accessibility as a proxy for potential protein-binding activity [31], we examined complementary ATAC-sequencing libraries [10] of the EndoC-βH1 untreated and cytokine-treated sample groups, once again focusing on RTEs. Consistent with what we observed for the transcriptome analysis, we observed a significant gain of accessibility at RTE loci upon proinflammatory cytokine stress (**Fig. 2a**). To examine the potential for specific TFs to bind to these loci, we utilized HOMER [32] to perform motif scanning (see Methods for details) and examine the enrichment of transcription factors. The top-most enriched TFs were from the interferon regulatory factor (IRF) family of TFs, which also exhibited the most dramatic increase in expression upon cytokine treatment (**Fig. 2b**). This pronounced enrichment suggests a prominent role of IRFs in the cytokine-induced activation of RTEs, contributing to the dysregulation of these elements and their subsequent effects. In addition, we decided to see what RTE subfamilies were most enriched and represented in the gain-of-accessibility loci. We observed LINE, SINE, and ERV subfamilies enriched at gain-of-accessibility loci (**Fig. 2c**), with the highest copy numbers originating from the L1 and *Alu* subfamilies (**Fig. 2d**). Footprinting analysis (see Methods for details) revealed signals for binding of inflammatory transcription factors, including IRFs, with increased signal in cytokine treated cells (**Fig. 2e** and **ESM** **Fig. 4**). Intriguingly, the highest enriched element by many orders of magnitude was the relatively unstudied MER107 element. An example of increased chromatin accessibility under cytokine treatment at an intergenic MER107 element (near a FLAM_C *Alu* element with a similarly prominent gain-of-accessibility peak) is shown in **Fig. 2e**. Taken together, these analyses indicate that proinflammatory stress treatment of human beta-cells can result in the dysregulation of RTEs capable of producing dsRNAs yielding deleterious downstream consequences.

**Fig. 4.**
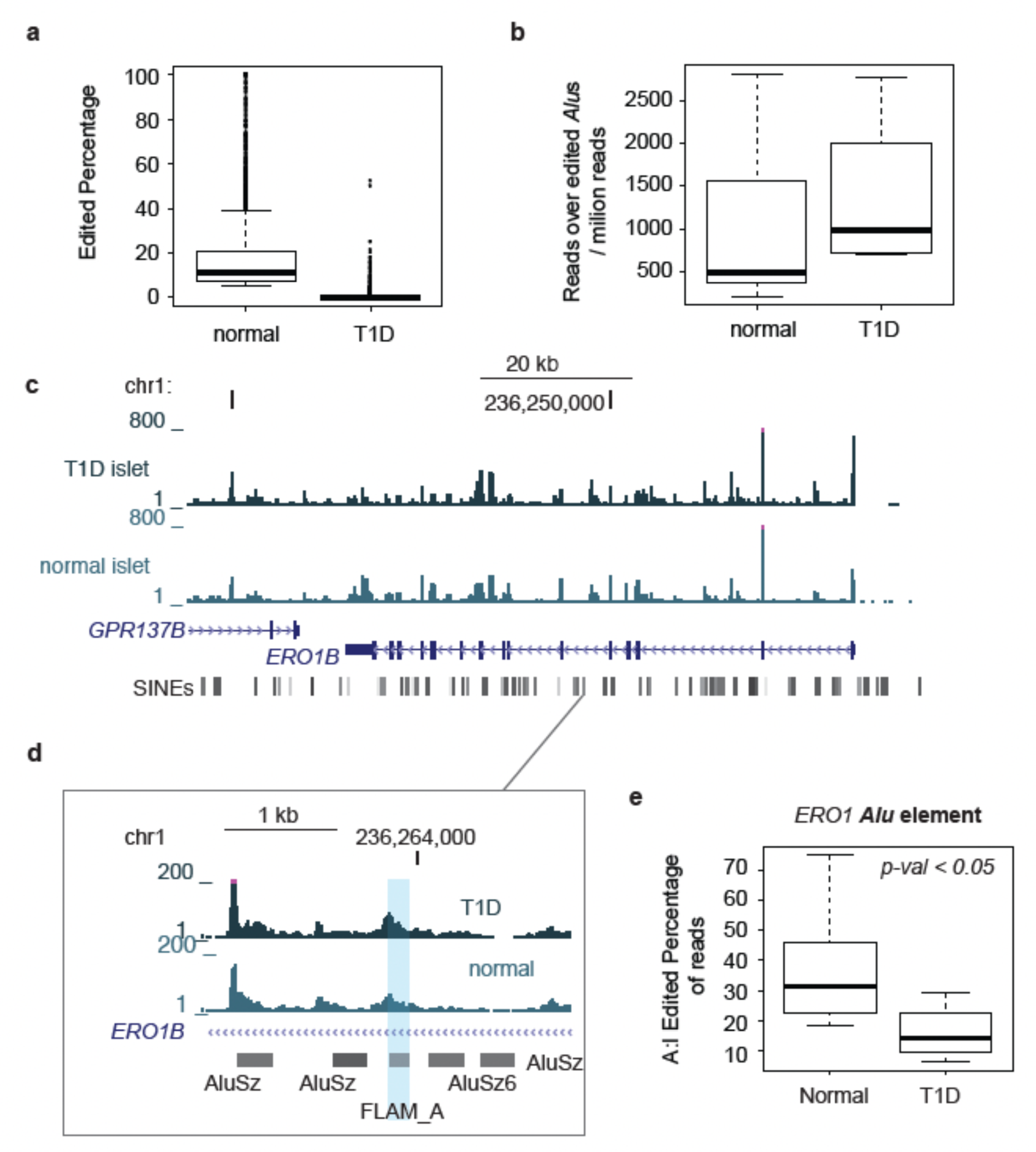
Islets from type 1 diabetics exhibit less A-to-I editing at *Alu* elements. (**a**) Comparison of the normalized number of reads found at *Alu*s with evidence of editing in normal (n = 12) and type 1 diabetic islets (n = 4). (**b**) Editing percentage of *Alu*s in both sample groups. (**c-e**) A genome-browser view of a locus with several representative *Alu*s. (**c**) RNA-sequencing data from normal and type 1 diabetic islets. (**d**) A zoomed-in view of *Alu*s of interest located within an intron of the ERO1B gene, showing that read number is similar between the two samples. (**e**) Quantification and comparison of A-to-I editing at the highlighted *Alu* in both sample groups (*p* < 0.05; Welch’s t-test).

### Unedited IR-Alus burden increases upon proinflammatory cytokine stress in human beta-cells

The ubiquitous nature of dsRNA structures circulating within the transcriptional pool of the cell – combined with their potential to push the cell into a destructive antiviral state – is almost exclusively resolved through A-to-I editing, catalyzed by ADAR1 [27, 28, 33, 34]. As previously mentioned, a large majority of these edited dsRNAs are inverted repeat *Alu*s (IR-*Alu*s) [5, 33]. In our next step, we wanted to examine the A-to-I editing levels of these *Alu*s in beta-cells. To gauge the extent of A-to-I editing, we analyzed the RNA-sequencing data with the *Alu* Editing Indexer (AEI) tool [33], which quantifies instances of A-to-I editing at annotated *Alu* loci (see Methods for details). Because A-to-I editing occurs only at dsRNAs, an advantage of using the AEI tool to examine editing events is that it also provides an implicit way to find *Alu*s with double-stranded nature.

We first filtered for and plotted up-regulated vs. down-regulated *Alu*s upon cytokine treatment. In similar fashion to the RTEs in general, *Alu*s were also significantly more up-regulated than down-regulated (**Fig. 3a**). We selected all up-regulated *Alu*s (see Methods for details) and then utilized AEI to determine the editing percentage. These analyses revealed that cytokine treatment increased the editing (**Fig. 3b**). This increase in editing percentage coincides with a significant increase in ADAR1 expression upon cytokine treatment (**Fig. 3c**). Though an increase in A-to-I editing would intuitively lead to a more ‘protective’ cellular environment against IR-*Alu* dsRNAs, evaluating the percentage of edited *Alu*s does not capture the total abundance of edited/unedited transcripts as the *pool* of *Alus* was larger in the cytokine-treated samples than in the control samples, as evidenced in the volcano plot (**Fig. 3a**). To account for this, we reanalyzed the RNA-seq data to count both the unedited and edited reads at each *Alu* locus within both the control and cytokine-treated sample groups. Instead of collapsing the editing data into a percentage, we scored each *Alu* locus on whether it was ‘more edited’ or ‘more unedited’ upon cytokine treatment (see Methods for details). Strikingly, we observed that, on average, there was a vastly significant higher amount of unedited IR-*Alu*s upon cytokine treatment despite a higher editing percentage (**Fig. 3d**). Thus, while the fraction of edited *Alu*s has increased, the overall abundance of unedited *<u>Alu</u>*s has also increased. Altogether, these results demonstrate that proinflammatory stress leads to an increased load of unedited IR-*Alu*s within human beta-cells.

### Beta-cells from type 1 diabetic individuals exhibit less A-to-I editing at IR-Alus when compared to non-diabetic beta-cells

We next examined whether A-to-I editing is altered in type 1 diabetic beta-cells versus non-diabetic beta-cells. We applied the *Alu* Editing Indexer to publicly available RNA-sequencing data [35] for cell-sorted beta-cells from either non-diabetic or diabetic patients to find evidence for viral mimicry acting *in vivo*. We found that *Alu*s with evidence of editing had increased read numbers in in beta-cells from type 1 diabetic individuals compared to non-diabetic cells (**Fig 4a**). There was, however, a notable decrease in the level of editing (∼1%) when compared to non-diabetic beta-cells (∼15%) (**Fig. 4b**). In similar fashion to the EndoC-βH1 cells, this discrepancy in editing was not a result of transcriptional change at *Alu*s – in fact, the observed higher read coverage of *Alu* loci in type 1 diabetic beta-cells bolsters the observation that more *Alu*s are expressed and less IR-*Alu* dsRNA transcripts are edited in the beta-cells from type 1 diabetic individuals (**Fig. 4a,e**). As an example, *ERO1B,* which is most prominently expressed in the pancreas, contains several *Alu* elements with less editing in type 1 diabetes patients than non-diabetic counterparts (**Fig. 4c,d**). Beta-cells from type 1 diabetics had significantly lower editing at this locus (∼15%) when compared to non-diabetic beta-cells (∼31%) (**Fig. 4e**). This decrease in A-to-I editing within beta-cells from type 1 diabetics suggests that their dsRNA pool is much more susceptible for detection by RIG-like receptors than the more edited dsRNA pool present in non-diabetic beta-cells. In conjunction with the inflammatory stress that drives the beta-cell into a much more vulnerable dsRNA-induced antiviral state, this impairment of A-to-I editing may further exacerbate the stress of the cell, accelerating it towards an IFN-I state.

## DISCUSSION

Our results introduce a model where the proinflammatory stress associated with insulitis primes the beta-cell towards a vulnerable and antiviral state through the emergence of endogenous unedited dsRNAs (IR-*Alu*s) within the cell. Current models of the events and mechanisms that lead to a full-scale adaptive immune cell-mediated attack on the beta-cells often cite defects in the immune system that drive attacks on otherwise normal beta-cells [36, 37]. Models also exist that describe the role of viral infections (most notably, Coxsackie B virus) as triggers to a more massive immune infiltration of the pancreatic islet [38, 39]. However, our results show that sterile (virus-free) stress conditions may lead to defects within the beta-cell, which prime the cell towards dysfunction and/or death. Furthermore, these defects are manifested through dsRNAs that arise from retroelement-derived loci such as IR-*Alu*s and ERVs, which contribute to a ‘viral mimicry’ phenotype.

The notion that stressors incite a dysregulation and misexpression of TEs has precedent – past studies have examined the loss of LINE-1 silencing during cellular senescence, causing them to become transcriptionally active, manifest as ssDNA in the cytosol, and drive an IFN-I response [40, 41]. Senescent beta-cells accumulated during autoimmune diabetes onset and progression are dysfunctional and exhibit an inflammatory paracrine phenotype, secreting damaging factors to healthy neighboring beta-cells that contribute to collateral beta-cell dysfunction/damage [41]. However, the connection between the inflammatory stress present during either beta-cell senescence or insulitis and the effect on the dysregulation of RTEs within the beta-cell is an area still unstudied. Our results show that inflammatory cytokine stress increases transcription at RTEs that manifest themselves as deleterious dsRNAs within the cell. Whether other stressors experienced by the beta-cells (e.g. ER stress, metabolic stress, etc.) can bring about a similar dysregulation of RTEs – and furthermore, whether such stressors cause a similar increase in dsRNAs within the beta-cell are still open questions.

The significant increase of dsRNAs we observed in islets upon inflammatory cytokine stress suggests that dsRNAs are involved in driving the IFN-I response. MDA5, one of the prominent RIG-like receptors for sensing viral dsRNAs within the cell, and its variants have gathered intrigue around its implication in the development of type 1 diabetes [11, 22, 42]. Additionally, the exploitation of the ‘viral mimicry’ phenomenon as a cancer therapeutic is primarily driven by chemical induction of RTE-derived dsRNAs within the cancer cells, leading to a deleterious IFN-I response driven through the MDA5/MAVS axis [1–3]. We show that upon proinflammatory cytokine exposure, human beta-cells are primed into an antiviral IFN-I state, characterized by (but not limited to) the significant upregulation of RLRs such as MDA5. These observations suggest that a *pathological-induction* (versus a pharmaceutical-induction) of RTE-derived dsRNAs within the cell – through inflammatory stress – may be an avenue in which beta-cells become defective in a virus-free/sterile microenvironment.

RTEs have been implicated in the progression of other autoimmune conditions such as multiple sclerosis, rheumatoid arthritis, and systemic lupus erythematosus [43–45]. Research surrounding the role of RTEs in these autoimmune conditions has largely centered around aberrant neoantigens produced from the endogenous retrovirus (ERV) subset of RTEs, such as HERV-W, HERV-K, and HERV-H [1–3]. Our work shown here pushes the role of RTEs as not just translation hotspots for neoantigens, but as transcriptional hotspots for dsRNAs. Additionally, our investigation of RTEs (most strikingly IR-*Alu*s) as deleterious species of dsRNAs has established a precedent in the field of type 1 diabetes pathogenesis. Our hypothesis that dsRNAs may be one avenue in how autoimmune diabetes can develop/exacerbate is further bolstered by the RNA-seq data showing a drastic decrease of A-to-I editing in the type 1 diabetic beta-cells compared to non-type 1 diabetic patients. This impairment in the dsRNA-editing capability of type 1 diabetic beta-cells lends further support to the deleterious nature of dsRNAs in such patients.

Altogether, our results show a novel way in which beta-cell defects can arise in an inflammatory cytokine microenvironment: through RTE-derived dsRNAs that are transcribed and uncover additional impacts that pro-inflammatory cytokines have on the human beta-cell. Furthermore, the capability to maintain this balance between autoimmunity and dsRNA-antiviral-immunity is diminished in type 1 diabetic beta-cells, suggesting that the beta-cells in diabetics are even more susceptible to be driven toward a viral mimicry IFN-I response, causing a collateral effect to neighboring beta-cells that may either render them dysfunctional or direct them towards death. With the existence of RIG-1 and reverse transcriptase inhibitors, these results have intriguing therapeutic implications as well.

## METHODS

### Cell culture, maintenance, and cytokine treatment

Human islets were obtained from the Integrated Islet Distribution Program (IIDP) at City of Hope. Collagenase-isolated human islets were hand-picked and sorted into 6-well plates and rested for 24 h in incubation media. Hu1168: age = 30, sex = female, HbA1c = 4.5%, no history of diabetes, CoD = anoxia/cardiovascular failure, purity = 85%, viability = 95%. Islets were then removed from incubation media and maintained and treated in CMRL media supplemented with 10% FBS, antibiotics, glutamine, beta-mercaptoethanol (50uM), and buffered with HEPES. IFNγ (100 ng/mL), TNF-α (10 ng/uL), and IL-1β (2 ng/uL) treatment of human islets was done for 24 h in CMRL growth media. 1.1B4 cells were generously donated from the Debbie Thurmond Lab (DMRI, City of Hope). 1.1B4 cells were kept, maintained, and passaged in DMEM media supplemented with 10% FBS, penicillin-strepomycin antibiotics, and glutamine. IFN-γ (100 ng/uL, PeproTech, CN: 300-02), TNF-α (10 ng/uL, PeproTech, CN: 300-01A), and IL-1β (2 ng/uL, PeproTech, CN: 200-01B) treatment of 1.1B4 cells was done for 24h in DMEM growth media. MIN6 cells were generously donated from the Ben Shih Lab (DMRI, City of Hope). MIN6 cells were kept, maintained, and passaged in RPMI-1640 media (low glucose, < 5mM) supplemented with 10% FBS, penicillin-streptomycin antibiotics, and glutamine. IFN-γ (100 ng/uL), TNF-α (10 ng/uL), and IL-1β (2 ng/uL) treatment of MIN6 was done for 24 h in RPMI-1630 media.

### Immunofluorescence microscopy and image processing

Treated or untreated human islets were fixed in 4% paraformaldehyde before being sent to the City of Hope Pathology Core for paraffin embedding and subsequent sectioning. Sections of human islets were de-paraffinized with xylene followed by serial washes of ethanol at decreasing concentrations, followed by rehydration/wash with PBS. Upon the de-paraffinization and rehydration of islet sections, sections were then blocked in blocking buffer (10% FCS, 3% BSA, 0.1% Triton-X in PBS) followed by incubation (1 hr) of α-dsRNA J2 primary antibody (1:100; SCICONS, U.S.). Cover slips were washed with PBST and incubated with secondary α-IgG AlexaFluor antibody (1:1000; Abcam) in blocking buffer (1 hr). To visualize nuclei, DAPI (10ug/mL in PBS) staining was done (5 min). α-Insulin antibody (1:200; Abcam) was also used to stain for beta-cells in the islet, followed by similar secondary antibody incubation. Immunofluorescence images were taken with a Zeiss LSM 700 Confocal Microscope. Images were processed and analyzed using Fiji/ImageJ to quantify signal of both dsRNA and insulin. Both 1.1B4 and MIN6 cells were split and grown upon circular cover lens within 24-well plates; MIN6 was seeded at 50,000 cells/well and MIN6 was seeded at 100,000 cells/well. Cells were washed (3x) with PBS and subsequently fixed to wells with 4% paraformaldehyde for 20 mins. Cells were permeabilized with PBST and blocked with blocking buffer (10% FCS, 3% BSA, 0.1% Triton-X in PBS). Cover slips were washed with PBST and incubated with secondary α-IgG AlexaFluor antibody (1:1000; Abcam) in blocking buffer (1 hr). To visualize nuclei, DAPI (10ug/mL in PBS) staining was done (5 min). To visualize insulin, α-Insulin antibody (1:200; Abcam) was used in similar incubation conditions. Immunofluorescence images were taken with a Zeiss LSM 700 Confocal Microscope, courtesy of City of Hope Microscopy Core. Images were processed and analyzed signal of both dsRNA and insulin.

### RNA-seq and ATAC-seq

EndoC-βH1 RNA-seq was obtained from GEO accession: GSE137136. EndoC-βH1 ATAC-seq was obtained from GEO accession: GSE123404. Type 1 diabetes beta-cell RNA-seq data was obtained from GEO accession GSE121863. RNA-seq data was trimmed with Trimgalore [46] and subsequently aligned uniquely to hg38 with STAR [47] v2.7.9a, using parameters –outMultimapperOrder Random, --outSAMmultNmax 1. Differential expression analysis of specific loci was done using DESeq2 [48]. GTF creation and transcription abundance analysis was performed using StringTie [49] v1.3.4d. ATAC-seq was trimmed with Trimgalore and subsequently aligned Bowtie2 [50] v2.4.1 to hg38, using parameters –very-sensitive, -k 10. Peak calling was performed with MACS2 [51] v2.2.5 callpeak, with parameters -p 0.05 --nomodel --shift -37 --extsize 73 --SPMR --keep-dup all --call-summits, to extract narrowPeak files. Differential analysis of peaks was performed with DESeq2 over repetitive element loci to give a set of gain-of-accessibility and loss-of-accessibility RTEs. Transcription factor analysis of ATAC-seq was performed with HOMER v4.10 [32] and TF motif overlaps were counted using the BedTools [52] suite. Volcano plots assess *p*-value with the Wald test and are corrected with the Benjamini-Hochberg method. Transcription factor footprinting was performed with HINT-ATAC [53], using motifs from JASPAR [54].

### A-to-I Editing Analysis

The *Alu* Editing Indexer (AEI) tool [33] was used to evaluate A-to-I editing instances over a given set of regions (e.g. *Alu*s). Common annotated SNPs are given to the tool to avoid misinterpretation of SNPs as editing events. BAM files were input into AEI to output .cmp files, which count the editing instances (by read coverage) within an *Alu* in single-nucleotide resolution. From the .cmp files, we calculated the editing percentage of an individual *Alu*:

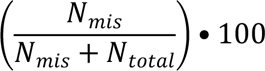

where *Nmis* is the number of A-to-G mismatches and *N*total is the total number of As in the length of the *Alu*. This percentage is taken across *Alu* loci output from the AEI across all samples, creating a comprehensive list. With the list of *Alu*s, count data is retrieved over each *Alu* from each sample and normalized, detailing the number of counts that each sample has over the specific *Alu* locus. With both the editing percentage and count amount taken from each of the *Alu* loci, we then multiplied them together to get the ‘edited counts’ fraction of the total counts. These ‘edited counts’ were subtracted from the ‘total counts’ to give us an ‘unedited counts’ number, representative of a percentage of unedited counts at a specific *Alu* locus.

## Supporting information

Electronic Supplementary Information

## Acknowledgements

Human islets were provided by the Integrated Islet Distribution Program (IIDP) and the Southern California Islet Cell Resource Center at City of Hope. We thank the City of Hope Pathology Core for helping with islet sectioning. We also thank members of the Schones lab that offered valuable feedback and revisions on this project.

## Data Availability

All raw and processed sequencing data generated in this study have been submitted to the NCBI Gene Expression Omnibus (GEO; https://www.ncbi.nlm.nih.gov/geo/) under accession number GSE247333.

## Authorship

K.C.: Contributed to writing of manuscript drafts, figure creation, experimental design/execution, conceptualization, and data analysis.

A.L.: Contributed to revision and writing of manuscript drafts, figure creation, and data analysis.

P.S.: Contributed to data analysis and figure creation.

D.S.: Contributed to writing/revision of manuscript drafts, figure creation, experimental conceptualization, and funding of project.

## Abbreviations

AEI: *Alu* Editing Indexer
ATAC-seq: Assay for Transposase-Accessible Chromatin sequencing
A-to-I: adenosine to inosine
GWAS: genome-wide association study
IFN-I: type 1 interferon
IFNB1: interferon-beta 1
dsRNA: double stranded RNA
IR-*Alu*s: inverted-repeat *Alu*s
RLRs: RIG-like receptors
RNA-seq: RNA-sequencing
RTE: retrotransposable element
TF: transcription factor.

## References

[1] Chiappinelli KB, Strissel PL, Desrichard A, et al. (2015) Inhibiting DNA Methylation Causes an Interferon Response in Cancer via dsRNA Including Endogenous Retroviruses. Cell 162(5): 974–986. 10.1016/j.cell.2015.07.011

[2] Roulois D, Loo Yau H, Singhania R, et al. (2015) DNA-Demethylating Agents Target Colorectal Cancer Cells by Inducing Viral Mimicry by Endogenous Transcripts. Cell 162(5): 961–973. 10.1016/j.cell.2015.07.056

[3] Liu M, Ohtani H, Zhou W, et al. (2016) Vitamin C increases viral mimicry induced by 5-aza-2’-deoxycytidine. Proc Natl Acad Sci U S A 113(37): 10238–10244. 10.1073/pnas.1612262113

[4] Mehdipour P, Marhon SA, Ettayebi I, et al. (2020) Epigenetic therapy induces transcription of inverted SINEs and ADAR1 dependency. Nature 588(7836): 169–173. 10.1038/s41586-020-2844-1

[5] Herzner AM, Khan Z, Van Nostrand EL, et al. (2021) ADAR and hnRNPC deficiency synergize in activating endogenous dsRNA-induced type I IFN responses. J Exp Med 218(9). 10.1084/jem.20201833

[6] Ahmad S, Mu X, Yang F, et al. (2018) Breaching Self-Tolerance to Alu Duplex RNA Underlies MDA5-Mediated Inflammation. Cell 172(4): 797–810 e713. 10.1016/j.cell.2017.12.016

[7] Campbell-Thompson M, Fu A, Kaddis JS, et al. (2016) Insulitis and beta-Cell Mass in the Natural History of Type 1 Diabetes. Diabetes 65(3): 719–731. 10.2337/db15-0779

[8] Nakamura T, Furuhashi M, Li P, et al. (2010) Double-stranded RNA-dependent protein kinase links pathogen sensing with stress and metabolic homeostasis. Cell 140(3): 338–348. 10.1016/j.cell.2010.01.001

[9] Newby BN, Mathews CE (2017) Type I Interferon Is a Catastrophic Feature of the Diabetic Islet Microenvironment. Front Endocrinol (Lausanne) 8: 232. 10.3389/fendo.2017.00232

[10] Ramos-Rodriguez M, Raurell-Vila H, Colli ML, et al. (2019) The impact of proinflammatory cytokines on the beta-cell regulatory landscape provides insights into the genetics of type 1 diabetes. Nat Genet 51(11): 1588–1595. 10.1038/s41588-019-0524-6

[11] Eizirik DL, Sammeth M, Bouckenooghe T, et al. (2012) The human pancreatic islet transcriptome: expression of candidate genes for type 1 diabetes and the impact of pro-inflammatory cytokines. PLoS Genet 8(3): e1002552. 10.1371/journal.pgen.1002552

[12] Colli ML, Ramos-Rodriguez M, Nakayasu ES, et al. (2020) An integrated multi-omics approach identifies the landscape of interferon-alpha-mediated responses of human pancreatic beta cells. Nat Commun 11(1): 2584. 10.1038/s41467-020-16327-0

[13] Eizirik DL, Colli ML, Ortis F (2009) The role of inflammation in insulitis and beta-cell loss in type 1 diabetes. Nat Rev Endocrinol 5(4): 219–226. 10.1038/nrendo.2009.21

[14] Avrahami D, Wang YJ, Klochendler A, Dor Y, Glaser B, Kaestner KH (2017) beta-Cells are not uniform after all-Novel insights into molecular heterogeneity of insulin-secreting cells. Diabetes Obes Metab 19 Suppl 1(Suppl 1): 147–152. 10.1111/dom.13019

[15] Chuong EB, Elde NC, Feschotte C (2017) Regulatory activities of transposable elements: from conflicts to benefits. Nat Rev Genet 18(2): 71–86. 10.1038/nrg.2016.139

[16] Chuong EB, Elde NC, Feschotte C (2016) Regulatory evolution of innate immunity through co-option of endogenous retroviruses. Science 351(6277): 1083–1087. 10.1126/science.aad5497

[17] Fueyo R, Judd J, Feschotte C, Wysocka J (2022) Roles of transposable elements in the regulation of mammalian transcription. Nat Rev Mol Cell Biol 23(7): 481–497. 10.1038/s41580-022-00457-y

[18] Dhillon P, Mulholland KA, Hu H, et al. (2023) Increased levels of endogenous retroviruses trigger fibroinflammation and play a role in kidney disease development. Nat Commun 14(1): 559. 10.1038/s41467-023-36212-w

[19] Rostami MR, Bradic M (2021) The derepression of transposable elements in lung cells is associated with the inflammatory response and gene activation in idiopathic pulmonary fibrosis. Mob DNA 12(1): 14. 10.1186/s13100-021-00241-3

[20] Paul DS, Teschendorff AE, Dang MA, et al. (2016) Increased DNA methylation variability in type 1 diabetes across three immune effector cell types. Nat Commun 7: 13555. 10.1038/ncomms13555

[21] Blum SI, Taylor JP, Barra JM, et al. (2023) MDA5-dependent responses contribute to autoimmune diabetes progression and hindrance. JCI Insight 8(2). 10.1172/jci.insight.157929

[22] Nejentsev S, Walker N, Riches D, Egholm M, Todd JA (2009) Rare variants of IFIH1, a gene implicated in antiviral responses, protect against type 1 diabetes. Science 324(5925): 387–389. 10.1126/science.1167728

[23] Smyth DJ, Cooper JD, Bailey R, et al. (2006) A genome-wide association study of nonsynonymous SNPs identifies a type 1 diabetes locus in the interferon-induced helicase (IFIH1) region. Nat Genet 38(6): 617–619. 10.1038/ng1800

[24] Lincez PJ, Shanina I, Horwitz MS (2015) Reduced expression of the MDA5 Gene IFIH1 prevents autoimmune diabetes. Diabetes 64(6): 2184–2193. 10.2337/db14-1223

[25] Lehmann KA, Bass BL (1999) The importance of internal loops within RNA substrates of ADAR1. J Mol Biol 291(1): 1–13. 10.1006/jmbi.1999.2914

[26] Pestal K, Funk CC, Snyder JM, Price ND, Treuting PM, Stetson DB (2015) Isoforms of RNA-Editing Enzyme ADAR1 Independently Control Nucleic Acid Sensor MDA5-Driven Autoimmunity and Multi-organ Development. Immunity 43(5): 933–944. 10.1016/j.immuni.2015.11.001

[27] Porath HT, Knisbacher BA, Eisenberg E, Levanon EY (2017) Massive A-to-I RNA editing is common across the Metazoa and correlates with dsRNA abundance. Genome Biol 18(1): 185. 10.1186/s13059-017-1315-y

[28] Slotkin W, Nishikura K (2013) Adenosine-to-inosine RNA editing and human disease. Genome Med 5(11): 105. 10.1186/gm508

[29] Heinrich MJ, Purcell CA, Pruijssers AJ, et al. (2019) Endogenous double-stranded Alu RNA elements stimulate IFN-responses in relapsing remitting multiple sclerosis. J Autoimmun 100: 40–51. 10.1016/j.jaut.2019.02.003

[30] Smit A, Hubley, R, Green, P. (2013-2015) RepeatMasker Open-4.0

[31] Tsompana M, Buck MJ (2014) Chromatin accessibility: a window into the genome. Epigenetics Chromatin 7(1): 33. 10.1186/1756-8935-7-33

[32] Heinz S, Benner C, Spann N, et al. (2010) Simple combinations of lineage-determining transcription factors prime cis-regulatory elements required for macrophage and B cell identities. Mol Cell 38(4): 576–589. 10.1016/j.molcel.2010.05.004

[33] Roth SH, Levanon EY, Eisenberg E (2019) Genome-wide quantification of ADAR adenosine-to-inosine RNA editing activity. Nat Methods 16(11): 1131–1138. 10.1038/s41592-019-0610-9

[34] Shadle SC, Bennett SR, Wong CJ, et al. (2019) DUX4-induced bidirectional HSATII satellite repeat transcripts form intranuclear double-stranded RNA foci in human cell models of FSHD. Hum Mol Genet 28(23): 3997–4011. 10.1093/hmg/ddz242

[35] Russell MA, Redick SD, Blodgett DM, et al. (2019) HLA Class II Antigen Processing and Presentation Pathway Components Demonstrated by Transcriptome and Protein Analyses of Islet beta-Cells From Donors With Type 1 Diabetes. Diabetes 68(5): 988–1001. 10.2337/db18-0686

[36] Knip M, Siljander H (2008) Autoimmune mechanisms in type 1 diabetes. Autoimmun Rev 7(7): 550–557. 10.1016/j.autrev.2008.04.008

[37] Roep BO, Peakman M (2012) Antigen targets of type 1 diabetes autoimmunity. Cold Spring Harb Perspect Med 2(4): a007781. 10.1101/cshperspect.a007781

[38] Skog O, Korsgren S, Melhus A, Korsgren O (2013) Revisiting the notion of type 1 diabetes being a T-cell-mediated autoimmune disease. Curr Opin Endocrinol Diabetes Obes 20(2): 118–123. 10.1097/MED.0b013e32835edb89

[39] Fairweather D, Rose NR (2002) Type 1 diabetes: virus infection or autoimmune disease? Nat Immunol 3(4): 338–340. 10.1038/ni0402-338

[40] Zhao K, Du J, Peng Y, et al. (2018) LINE1 contributes to autoimmunity through both RIG-I- and MDA5-mediated RNA sensing pathways. J Autoimmun 90: 105–115. 10.1016/j.jaut.2018.02.007

[41] De Cecco M, Ito T, Petrashen AP, et al. (2019) L1 drives IFN in senescent cells and promotes age-associated inflammation. Nature 566(7742): 73–78. 10.1038/s41586-018-0784-9

[42] Shigemoto T, Kageyama M, Hirai R, Zheng J, Yoneyama M, Fujita T (2009) Identification of loss of function mutations in human genes encoding RIG-I and MDA5: implications for resistance to type I diabetes. J Biol Chem 284(20): 13348–13354. 10.1074/jbc.M809449200

[43] Hansen MP, Matheis N, Kahaly GJ (2015) Type 1 diabetes and polyglandular autoimmune syndrome: A review. World J Diabetes 6(1): 67–79. 10.4239/wjd.v6.i1.67

[44] Triolo TM, Armstrong TK, McFann K, et al. (2011) Additional autoimmune disease found in 33% of patients at type 1 diabetes onset. Diabetes Care 34(5): 1211–1213. 10.2337/dc10-1756

[45] Nexo BA, Villesen P, Nissen KK, et al. (2016) Are human endogenous retroviruses triggers of autoimmune diseases? Unveiling associations of three diseases and viral loci. Immunol Res 64(1): 55–63. 10.1007/s12026-015-8671-z

[46] Martin M (2011) Cutadapt Removes Adapter Sequences from High-Throughput Sequencing Reads. EMBnet Journal 17. 10.14806/ej.17.1.200

[47] Dobin A, Davis CA, Schlesinger F, et al. (2013) STAR: ultrafast universal RNA-seq aligner. Bioinformatics 29(1): 15–21. 10.1093/bioinformatics/bts635

[48] Love MI, Huber W, Anders S (2014) Moderated estimation of fold change and dispersion for RNA-seq data with DESeq2. Genome Biol 15(12): 550. 10.1186/s13059-014-0550-8

[49] Pertea M, Pertea GM, Antonescu CM, Chang TC, Mendell JT, Salzberg SL (2015) StringTie enables improved reconstruction of a transcriptome from RNA-seq reads. Nat Biotechnol 33(3): 290–295. 10.1038/nbt.3122

[50] Langmead B, Salzberg SL (2012) Fast gapped-read alignment with Bowtie 2. Nat Methods 9(4): 357–359. 10.1038/nmeth.1923

[51] Zhang Y, Liu T, Meyer CA, et al. (2008) Model-based analysis of ChIP-Seq (MACS). Genome Biol 9(9): R137. 10.1186/gb-2008-9-9-r137

[52] Quinlan AR, Hall IM (2010) BEDTools: a flexible suite of utilities for comparing genomic features. Bioinformatics 26(6): 841–842. 10.1093/bioinformatics/btq033

[53] Li Z, Schulz MH, Look T, Begemann M, Zenke M, Costa IG (2019) Identification of transcription factor binding sites using ATAC-seq. Genome Biol 20(1): 45. 10.1186/s13059-019-1642-2

[54] Castro-Mondragon JA, Riudavets-Puig R, Rauluseviciute I, et al. (2022) JASPAR 2022: the 9th release of the open-access database of transcription factor binding profiles. Nucleic Acids Res 50(D1): D165–D173. 10.1093/nar/gkab1113

