## Supplementary material for "Stress-induced endogenous-dsRNAs drive beta-cells to IFN-I state": Electronic Supplementary Information

### Materials and Methods – Supplemental

*Transfection of MDA5*

pFLAG-CMV4-MDA5wt and pFLAG-CMV4-MDA5G495R were generously provided by the Sun Hur lab. MIN6 cells were transfected with Lipofectamine 3000 (Invitrogen, L3000001) (as per instructions provided by manufacturer) with either plasmid and allowed to grow for 2 d before collection of RNA for qPCR analysis.

*RNA-seq of 1.1B4 and MIN6 cells*

To prepare MIN6 and 1.1B4 RNA-seq, treated cells were washed with PBS, lysed with TRIzol reagent and incubated. Each sample group was done in triplicate (Ncontrol = 3, Ncyto = 3, for a total of 6 libraries per cell line). Hydrophilic layer was separated and subsequently extracted for RNA with Quick-RNa Miniprep Kit (Zymo Research). RNA library was constructed with the help of the City of Hope Integrative Genomics Core, depleted of rRNA with Invitrogen RiboMinus, and sequenced at a depth of 25 million reads/sample. Raw RNA-seq FASTQs were trimmed with Trimgalore [1] and subsequently aligned uniquely to hg38/mm10 (respectively) with STAR [2] v2.7.9a, using parameters –outMultimapperOrder Random, --outSAMmultNmax 1. Differential expression analysis of specific loci was performed using DESeq2 [3] and plots were created with ggplot2 package in R v4.3.1.

*Colony-forming/Clonogenic Assay*

1.1B4 cells were grown and treated accordingly for 24 h, with or without TLR3i

(CalBiochem, USA). Each sample group was then detached with trypsin and counted for initial seeding in 6-well plates and seeded at 150 cells/well. Cells were then given fresh media (with or without cytokine/TLR3i treatment) and allowed to grow for 1 week, with media being changed out after 72 hours. Cells were washed and fixed with 100% methanol and incubated for 20 mins. Cells were rinsed with water and then incubated with crystal violet solution for 5 mins to visualize colonies. CV was then washed away with water until excess dye was removed and colonies were then dried overnight. Plates were visualized with bright-field microscope and colonies were manually counted. Image processing was performed with Fiji/ImageJ.

*Electron Microscopy*

EM was performed as described in Ahmad et al. To summarize, cytosolic RNA (600 ug/ul) was extracted from untreated/cytokine-treated MIN6 cells. RNA was then preincubated with MDA5wt (450 nM) or in incubating buffer (20 mM HEPES, pH 7.5, 50 mM NaCl, 2 mM MgCl2, and 2 mM DTT) alone for 10 mins at room temperature, followed by addition of 1 mM ADP • AlFx on ice. The ratio of ADP • AlFx was prepared with ADP, AlCl3, and NaF in a molar ratio of 1:1:3. With the help of City of Hope EM Core, filaments were prepared with 0.75% uranyl formate and then examined under TEM microscopy.

### References


[1] Martin M (2011) Cutadapt Removes Adapter Sequences from High-Throughput Sequencing Reads. EMBnet Journal 17. https://doi.org/10.14806/ej.17.1.200

[2] Dobin A, Davis CA, Schlesinger F, et al. (2013) STAR: ultrafast universal RNA-seq aligner. Bioinformatics 29(1): 15-21. 10.1093/bioinformatics/bts635

[3] Love MI, Huber W, Anders S (2014) Moderated estimation of fold change and dispersion for RNA-seq data with DESeq2. Genome Biol 15(12): 550. 10.1186/s13059-014-0550-8

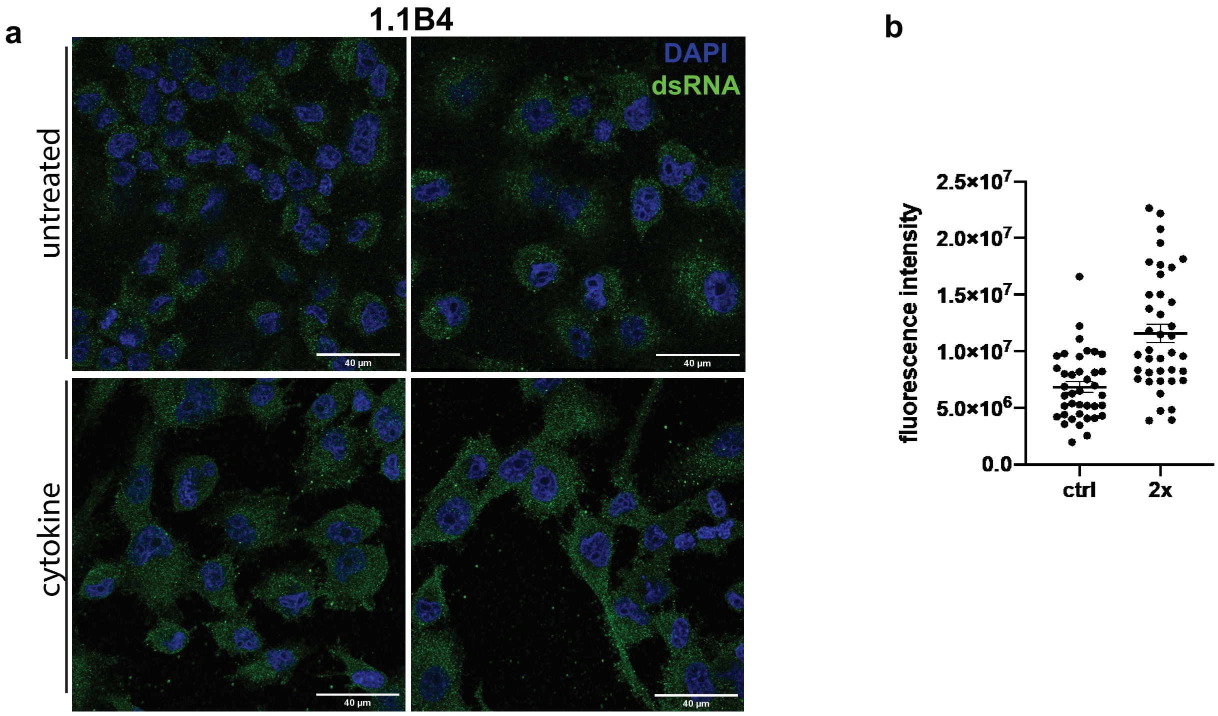


**ESM Fig. 1 - dsRNAs are increased in 1.1B4 cells upon proinflammatory cytokine stress**. 1.1B4 cells were treated with IFNg + TNFa for 24 h before staining and imaging.(a) Immunofluorescence of 1.1B4 staining for dsRNAs (green) and with DAPI staining for nuclei (blue). White scale bar represents 40um. (b) dsRNA fluorescence was normalized and quantified using Fiji/ImageJ

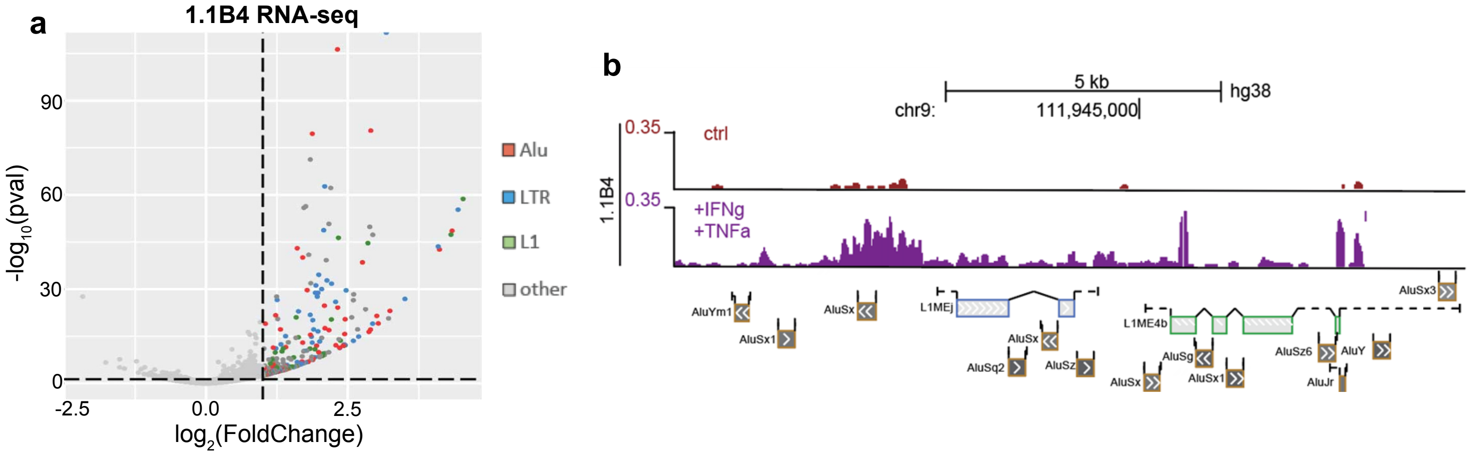


**ESM Fig. 2** - Transcription from RTE loci is upregulated upon cytokine treatment in 1.1B4 cells. (a) Volcano plot of RNA-sequencing from untreated (n = 3) vs. cytokine-treated (IFNg + TNFa) 1.1B4 cells (n = 3). Each point represents an annotated retrotransposon, as denoted by RepeatMasker (hg38). (b) Representative view of the RNA-seq signal at an example intergenic locus, showing the characteristics of cytokine-induced increase of expression at retrotransposons.

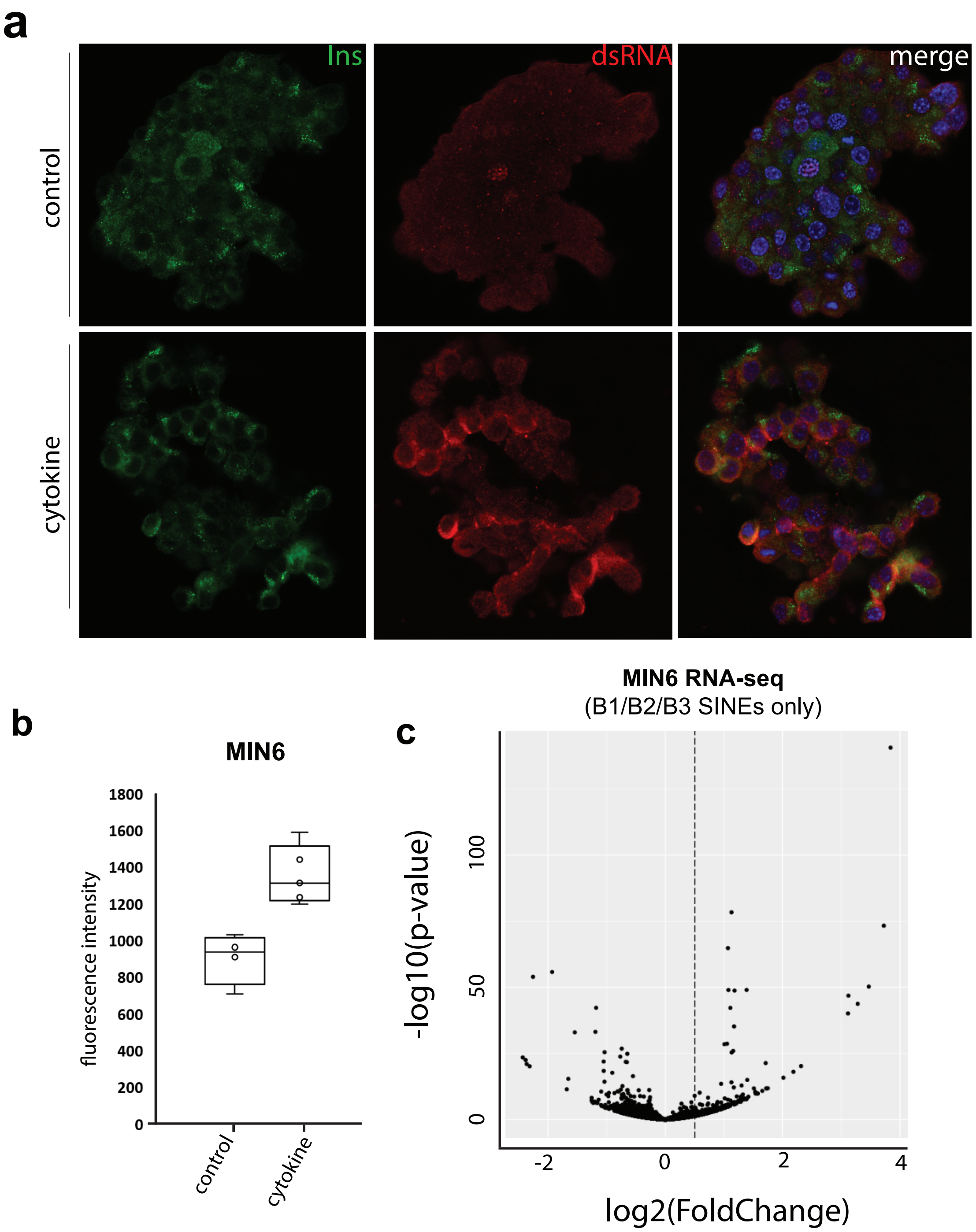


**ESM Fig. 3 - dsRNAs are increased in MIN6 cells upon proinflammatory cytokine stress, with upregulation of murine SINE elements.** MIN6 cells were treated with IFNg + TNFa for 24 h before staining and imaging. (a) Immunofluorescence of MIN6 staining for dsRNAs (red), and insulin (magenta), with DAPI staining for nuclei (blue). (b) dsRNA fluorescence was normalized and quantified using Fiji/ImageJ. (c) Volcano plot showing the transcriptional changes at B1/B2/B3 SINE loci within MIN6 upon cytokine treatment. MIN6 RNA-seq was performed on untreated (n = 3) and cytokine-treated (n= 3) samples.

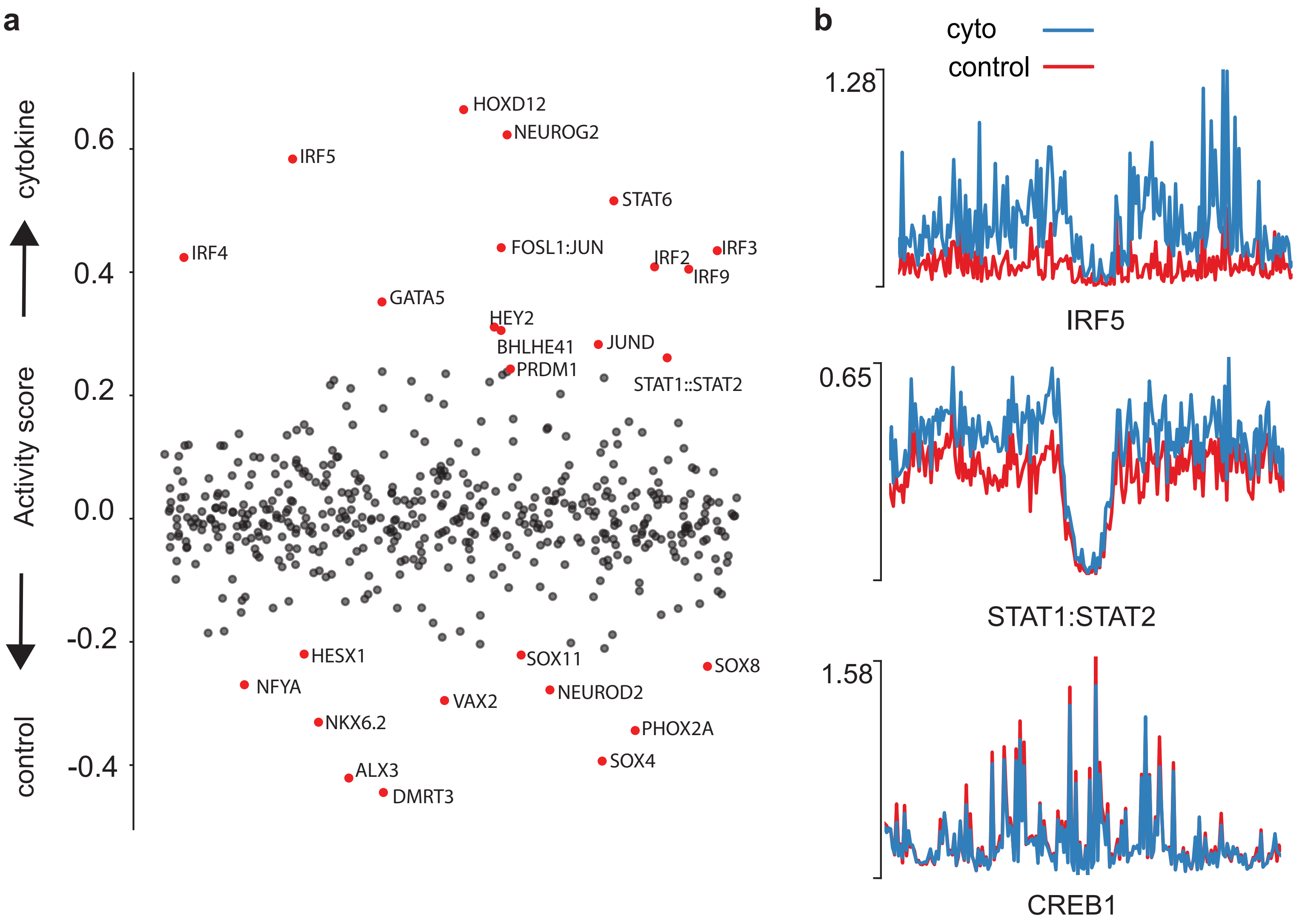


**ESM Fig. 4 - Edited Alus contain motifs for inflammation-associated transcription factors.** Footprinting analysis shows that edited Alus are enriched for transcription factors that emerge from insulitic stress. (a) Scatter plot showing the "activity score" of all JASPAR repository TFs. Activity score is an index that is a function of how well protected the footprint is and the 'openness' of the site which measures the number of cleavage events around the motif. Dots towards the top show TFs with more activity in cytokine-treated samples while dots toward the bottom show TFs with more activity in control-treated samples. (b) Footprint plots showing the bias-corrected normalized signal at the corresponding inflammatory TF motifs (IRF5, STAT1:STAT2 heterodimer) and CREB1 as a control. Red is control signal; blue is cytokine signal.

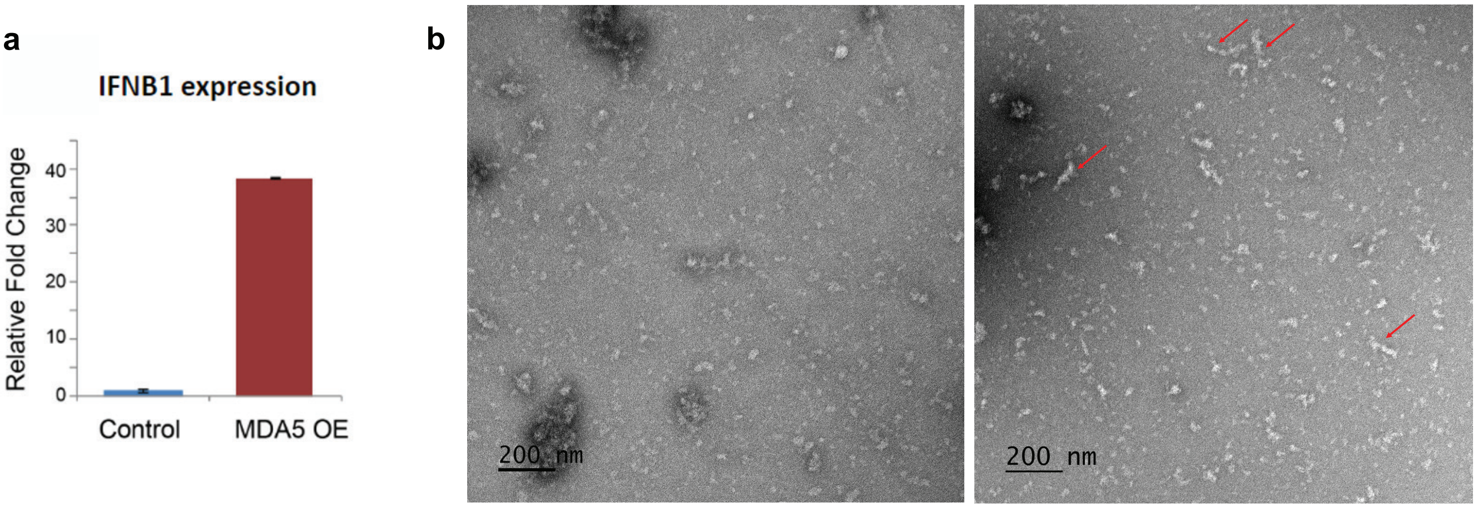


**ESM Fig. 5- Cytosolic RNA from cytokine-stress MIN6 cells exhibits more MDA5 filamentation.** (a) qPCR of IFNB1 expression in MIN6 cells upon either sham-shRNA or overexpression of wild-type MDA5. (b) TEM images of purified recombinant wild-type MDA5 in situ with cytosolic RNA fractions of either (left) control MIN6 cytoRNA or (right) cytokine-stressed cytoRNA.
